# Locating the ligand binding Sites for the G-protein coupled estrogen receptor (GPER) using combined information from docking and sequence conservation

**DOI:** 10.1101/061051

**Authors:** Ashley R. Vidad, Stephen Macaspac, Ho Leung Ng

## Abstract

High concentrations of estrogenic compounds can overstimulate estrogen receptors and potentially lead to breast, ovarian, and cervical cancers. Recently, a G-protein coupled estrogen receptor (GPER/GPR30) was discovered that has no structural similarity to the well-characterized, classical estrogen receptor ERα. The crystal structure of GPER has not yet been determined, and the ligand binding sites have not yet been experimentally identified. The recent explosion of GPCR crystal structures now allow homology modeling with unprecedented reliability. We create, validate, and describe a homology model for GPER. We describe and apply ConDock, the first hybrid scoring function to use information from protein surface conservation and ligand docking, to predict binding sites on GPER for four ligands, estradiol, G1, G15, and tamoxifen. ConDock is a simple product function of sequence conservation and binding energy scores. ConDock predicts that all four ligands bind to the same location on GPER, centered on L119, H307, and N310; this site is deeper in the receptor cleft than are ligand binding sites predicted by previous studies. We compare the sites predicted by ConDock and traditional methods analyzing surface geometry, surface conservation, and ligand chemical interactions. Incorporating sequence conservation information in ConDock avoids errors resulting from physics-based scoring functions and modeling.

## Introduction

Estrogens are hormones with multiple physiological and pathological functions in both men and women. The primary reproductive hormone in females, estrogen is responsible for ovarian development, oocyte and endometrial maturation, and uterine contraction^1^. Overexpression of estrogens is associated with estrogen-sensitive ovarian, endometrial, and breast cancers^2^. In addition to its role in the female reproductive system, estrogen is also an important regulator in the skeletal, nervous, cardiovascular, endocrine, and immune systems^3^. Estrogen functions in production of bone, insulin secretion, T cell differentiation, neuroprotection, and modulation of pain sensation, and cardiomyocyte contractility^3–5^. Imbalance of estrogen levels can lead to various pathological disorders such as osteoporosis, diabetes mellitus, autoimmune diseases, and dilated myopathy^3–6^.

The most important estrogen is 17β-estradiol (E2). The primary estrogen receptors, estrogen receptor-α (ERα) and estrogen receptor-β (ERβ) are dimeric nuclear receptor transcription factors that activate gene expression when bound to E2^7,8^. These two receptors are structurally very similar. Depending on tissue and ligand, ERα and ERβ can act redundantly, synergistically, and/or antagonistically. Studies suggest that ERα and ERβ have minimal participation in rapid and membrane-associated estrogen signaling activities as shown by the continued activation by estrogen in the presence of ER antagonists^9^. This suggested the existence of unidentified membrane-associated estrogen receptors. G-protein coupled estrogen receptor (GPER, formerly known as GPR30) is a membrane-bound ER that was discovered recently; GPER is proposed to be a primary mediator of rapid estrogen-associated effects^10–12^. It has also been shown to mediate other estrogen-activated responses such as cAMP regeneration and nerve growth factor expression^13,14^. Because of the technical challenges involved in experiments with G-protein coupled receptors, biochemical experiments are difficult and no crystal structure of GPER is available, and thus details of ligand binding are unknown.

Estrogen receptors can recognize a range of structurally diverse ligands (Fig. 1). E2 is the primary ligand of well-characterized estrogen receptors. E2 binds to all known estrogen receptors and is thus non-specific for GPER. Estrogen receptor binding ligands, such as tamoxifen, are also used as drugs to treat estrogen-responsive breast cancer^15,16^. Tamoxifen acts as an ER antagonist in some tissues and as an ER agonist in others, which is why it is considered a selective estrogen receptor modulator (SERM)^17^. Other drugs such as fulvestrant act as ER antagonists in all tissues; these drugs are considered to be selective estrogen receptor downregulators (SERD)^18^. Some SERMs and SERDs are agonists of GPER^9^, and recently GPER-specific ligands G1 and G15 were discovered^19,20^. G1 and G15 are structurally similar; they differ by an acetyl group (Fig. 1). G1 is an agonist, whereas G15 is an antagonist.

**Figure 1.**
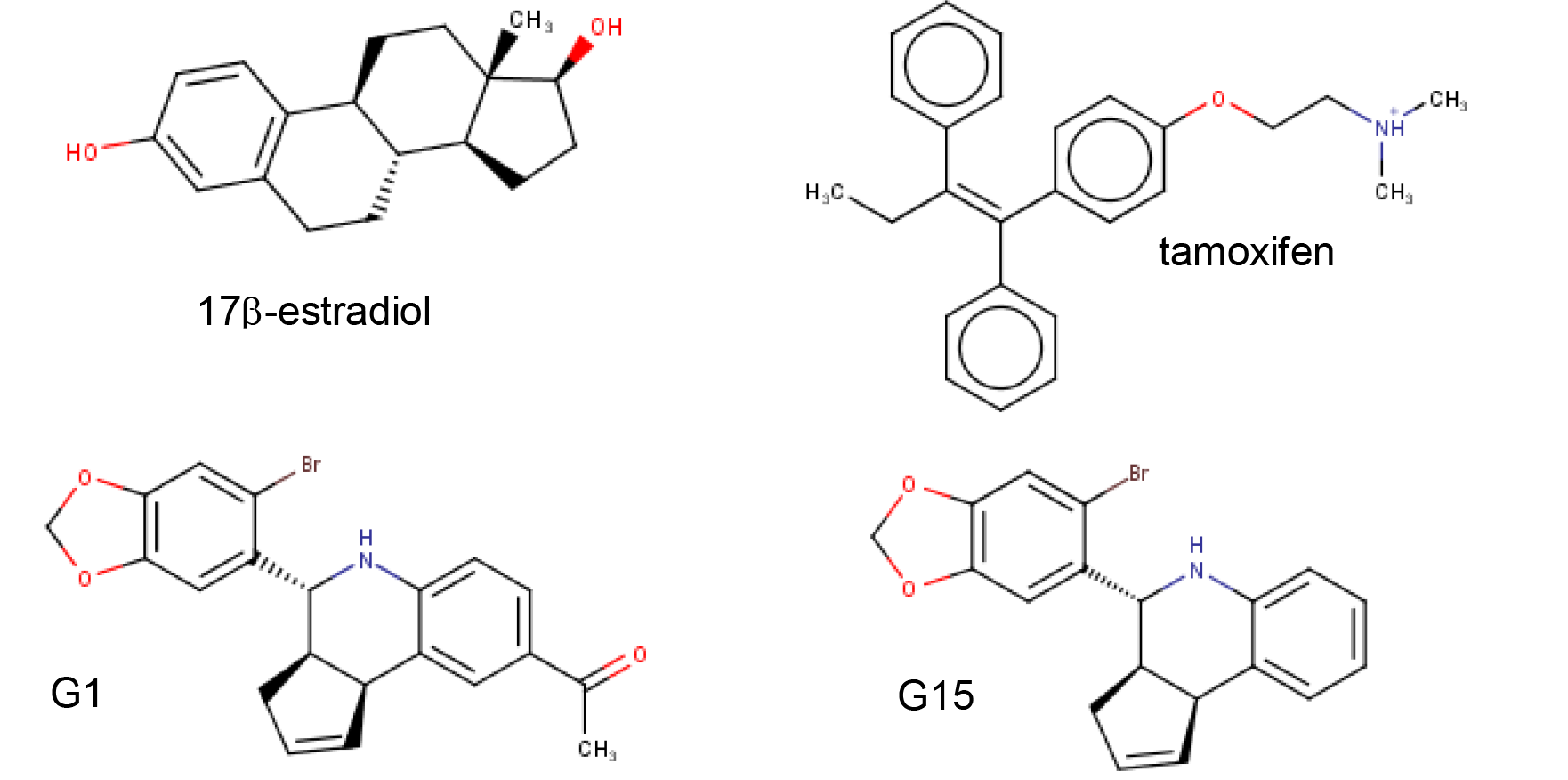
Four experimentally verified GPER ligands

The ligand binding sites in GPER have not yet been experimentally identified. Various computational approaches have been used to predict ligand binding sites in other G-protein coupled receptors. Traditional docking methods compute the lowest energy orientation of a ligand fit to a receptor surface. Such methods are highly dependent on the form of the energy scoring function and accuracy of the receptor model structure^21–23^. These methods have been used to identify ligand binding sites and build pharmacophores for G-protein coupled receptors (GPCRs)^24–26^, but the lack of diverse GPCR crystal structures presents serious challenges to using docking methods for identification of ligand binding sites.

Another feature that can be used to predict ligand binding sites is surface or sequence conservation. Binding sites for particular ligands are often conserved, although systematic sequence variation can encode ligand specificity^27–29^. The massive abundance of genomic data for GPCRs strongly constrains possibilities for ligand binding sites even without chemical or structural information^30–32^.

There has been less research on methods that combine information from chemical interactions, geometric surface analysis, and bioinformatics. Hybrid strategies, such as Concavity^33^, have demonstrated superior performance in predicting ligand binding sites compared to single-mode approaches. Concavity scores binding sites by evolutionary sequence conservation as quantified by the Jensen-Shannon divergence^29^ and employs geometric criteria of size and shape. Here, we describe and apply a new hybrid scoring function, ConDock, which combines information from surface conservation with intermolecular interactions from docking calculations, to predict ligand binding sites for GPER. We compare our results from those previously published using purely docking based methods^34,35^.

## Results

Because there are no crystal structures yet available for GPER, we created a homology model using GPCR-I-TASSER^36^, among the most reliable GPCR homology modeling software packages. GPCR-I-TASSER identified the closest matching crystal structure to GPER as that of the CCR5 chemokine receptor (PDB 4mbs) with 23% sequence identity. GPCR-I-TASSER used this crystal structure along with 9 other GPCR crystal structures as templates for homology modeling. The GPER homology model differs from chain A of the crystal structure of CCR5 chemokine receptor with RMSD of 0.96 Å across Cα atoms (Fig. S1). The primary differences are in the extracellular loop between helices 4 and 5 and the intracellular loops between helices 5 and 6, and after helix 7. These two intracellular loops are those predicted by ERRAT^37^ to be most likely in error based on the likelihood of atom pair type interactions from high resolution crystal structures. It should be noted that ERRAT was not developed for analyzing membrane proteins so the reliability of its error analysis for the GPER homology model is uncertain. Furthermore, as the sites with the greatest predicted errors are on the intracellular face of GPER, far from the ligand binding site, they are less likely to affect our ligand prediction study.

Using the SwissDock server^38^, we docked structures of the four ligands E2, G1, G15, and tamoxifen (Fig. 1) to a homology model of GPER, as no crystal structure is available. Most of the docked sites from SwissDock were not located on the extracellular face of GPER and thus were considered nonviable (Fig. S3). The shortcomings of a purely physics based scoring function such as that used by SwissDock in predicting ligand binding is not surprising given the lack of an experimental crystal structure and well known limitations of current computational methodology^39–42^.

We next ranked all ligand binding sites generated by SwissDock that were located on the extracellular face of GPER using the combined ConDock score. The ConDock score is the simple product of the ConSurf^43,44^ binding surface sequence conservation score and the SwissDock FullFitness energy score^45^. A highly negative ConDock score is associated with a more probable ligand binding site. For all four ligands, the ConDock score identified one or two ligand binding sites and poses that clearly outscored other candidates (Table 1). ConDock identified the same approximate binding site for all four ligands, although this was not an explicit criterion in the calculations (Fig. 2). The average ConSurf conservation score across the four ligand binding sites is 0.82, indicating that the site is highly but not completely conserved. The binding site is located deep in the receptor cleft, although this too was not a criterion in the prediction calculation. Given the lack of additional experimental evidence for the location of the ligand binding site, the proposed ConDock sites are physically reasonable.

**Table 1.**
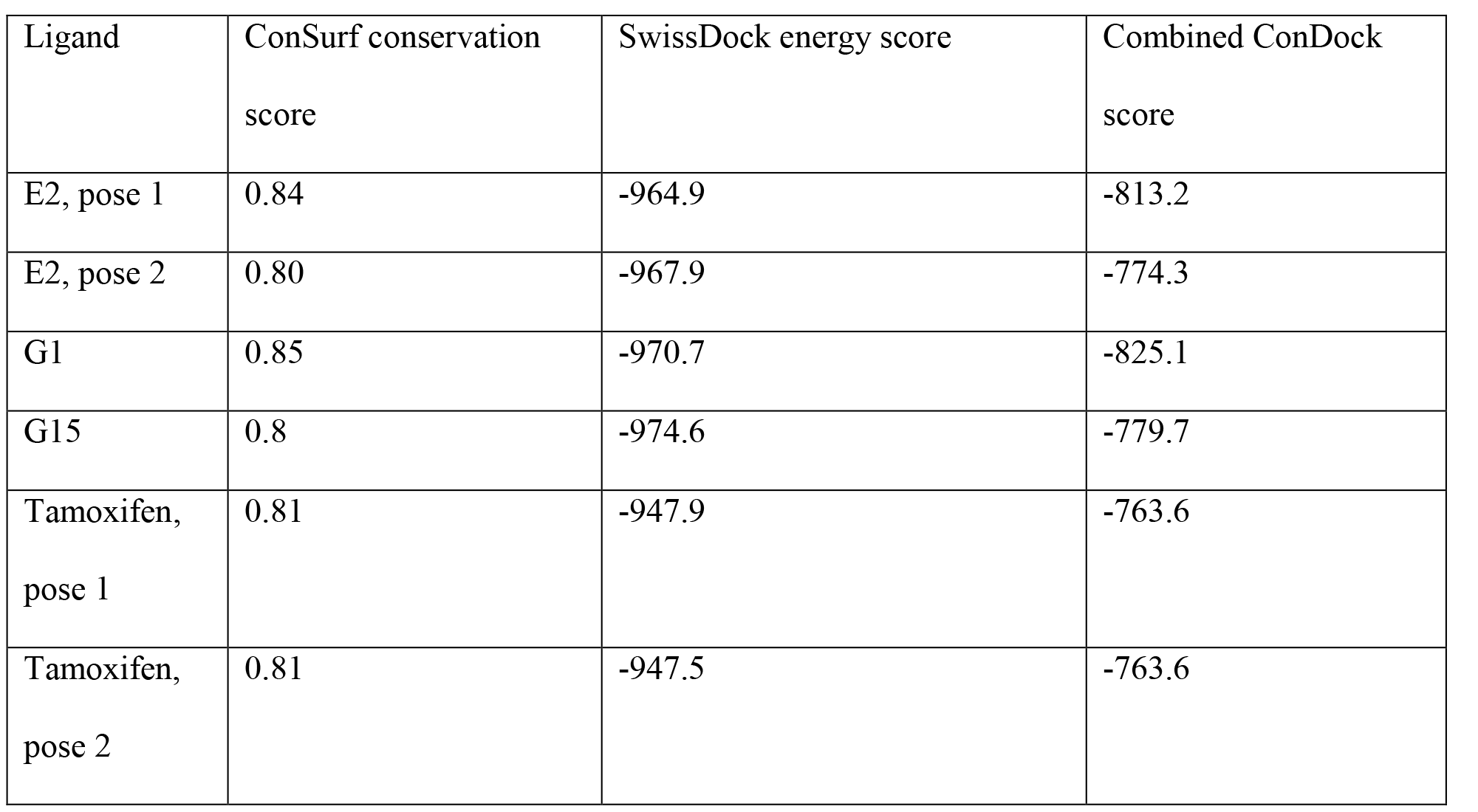
Scores for predicted binding sites and poses for GPER ligands.

**Figure 2.**
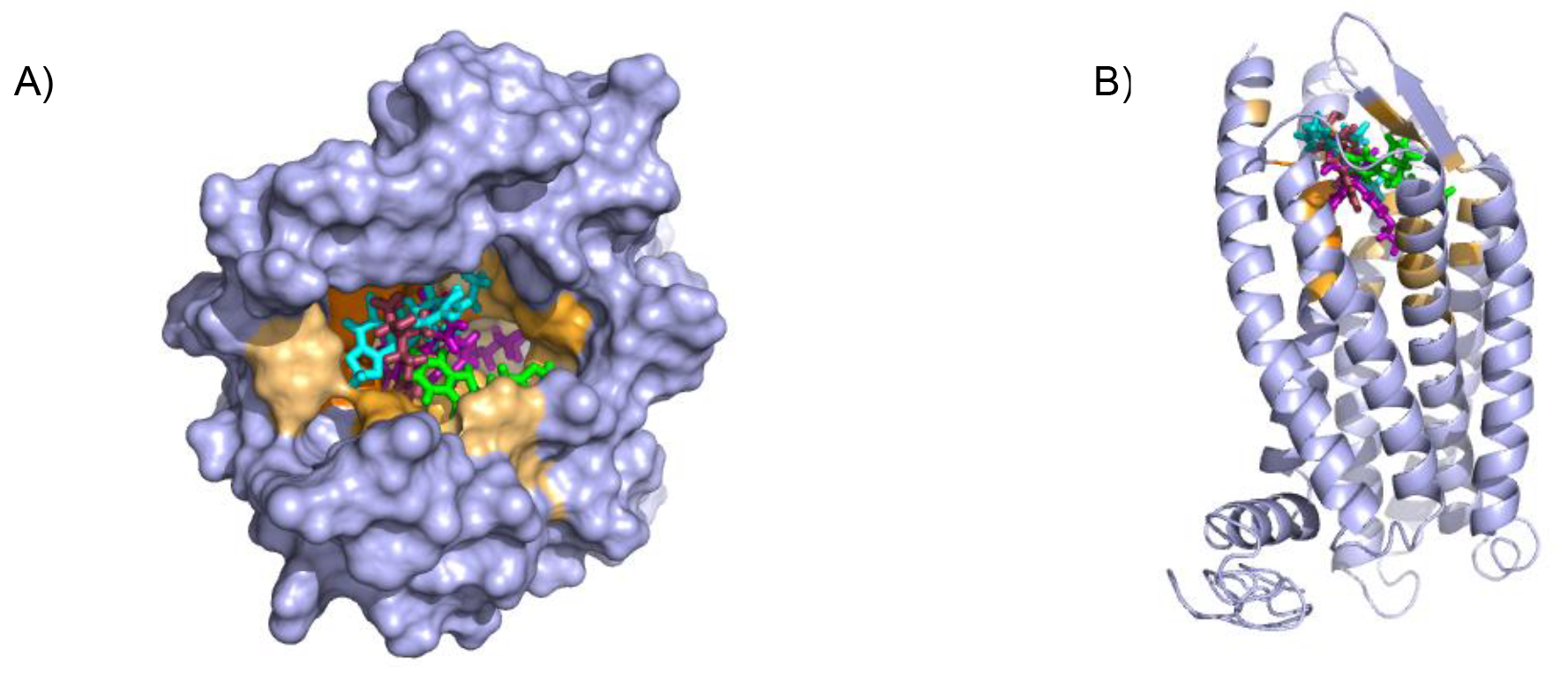
Predicted ligand binding pockets in GPER. A) Extracellular perspective of GPER showing amino acids predicted to contact ligands E2 (maroon), G1 (cyan), G15 (green), and tamoxifen (violet). Residues colored orange are predicted to contact one or more ligands with darker hue indicating interaction with multiple ligands. B) Predicted binding pocket viewed from a 90° rotation.

We found two promising binding sites for E2 in GPER. The two sites are 4.4 Å apart, located deep in the receptor cleft (Fig. 3). E2 is oriented perpendicular to the lipid membrane and rotated about 180° between the two poses. The conservation scores for these two poses are 0.84 and 0.80. The energy scores of the two poses are similar. The amino acids contacting E2 in pose 1 are conserved in GPERs from six species, and only one residue contacting pose 2, H282, varies across species. In the top ranked pose, there is a hydrogen bond between the inward pointing D-ring hydroxyl group of E2 and the carboxyl terminal on E115. Hydrophobic interactions are present between E2 and non-polar residues L119, Y123, P303, and F314. In the second ranked pose, the inward pointing A-ring hydroxyl group of E2 makes a hydrogen bond with N310. This pose is in a less hydrophobic environment, contacting primarily H282 and P303.

**Figure 3.**
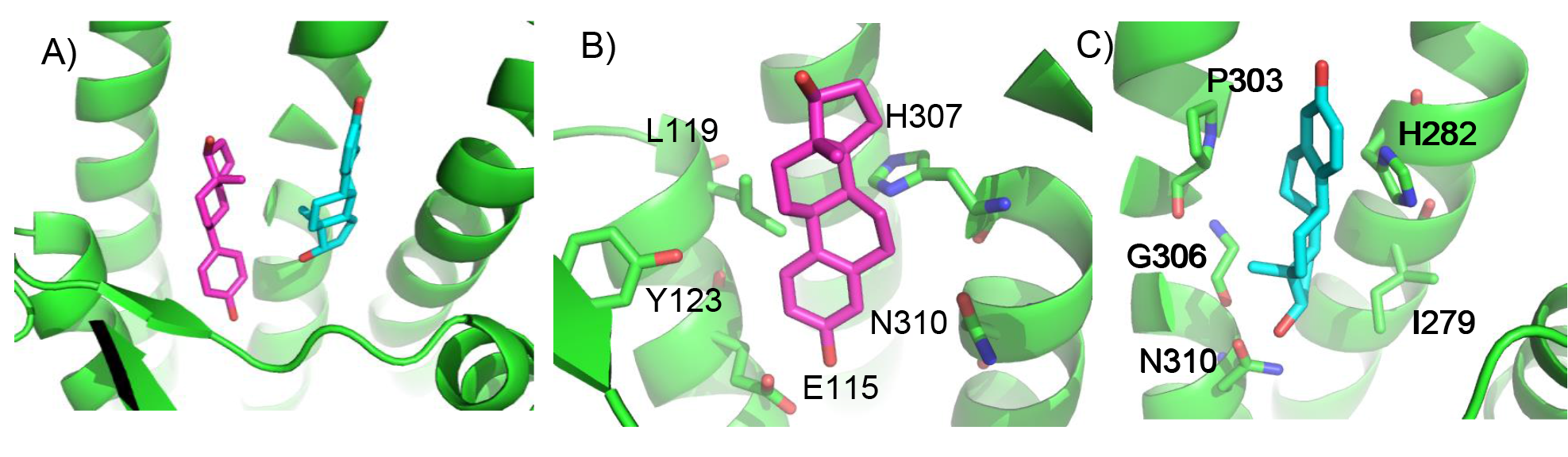
Predicted E2 binding sites in GPER. A) The two highest scoring docking poses for E2. B) Receptor-ligand interactions for E2 pose 1. C) Receptor-ligand interactions for E2 pose 2.

ConDock predicts that G1 and G15 bind in adjacent but distinct binding sites separated by 2.3 Å despite the chemical similarity of the two ligands. The top predicted binding site for G1 is found within the pocket bound by Y55, L119, F206, Q215, I279, P303, H307, and N310 (Fig. 4). This orientation had the highest conservation score of all predicted binding sites at 0.85. In this pose, N310 makes a long hydrogen bond with the acetyl oxygen of G1. The predicted binding site for G15 is found within the pocket bound by L119, Y123, M133, S134, L137, Q138, P192, V196, F206, C207, F208, A209, V214, E218, H307, and N310. This pose had a conservation score of 0.8. Hydrogen bonding is not observed between GPER and G15. Hydrophobic interactions are observed with L119, Y123, F206, and V214. An interesting aspect of this binding site is that the amino acid at position 214 is valine in humans but is isoleucine in Atlantic croaker and zebrafish. These two hydrophobic amino acids differ by only a methyl group.

**Figure 4.**
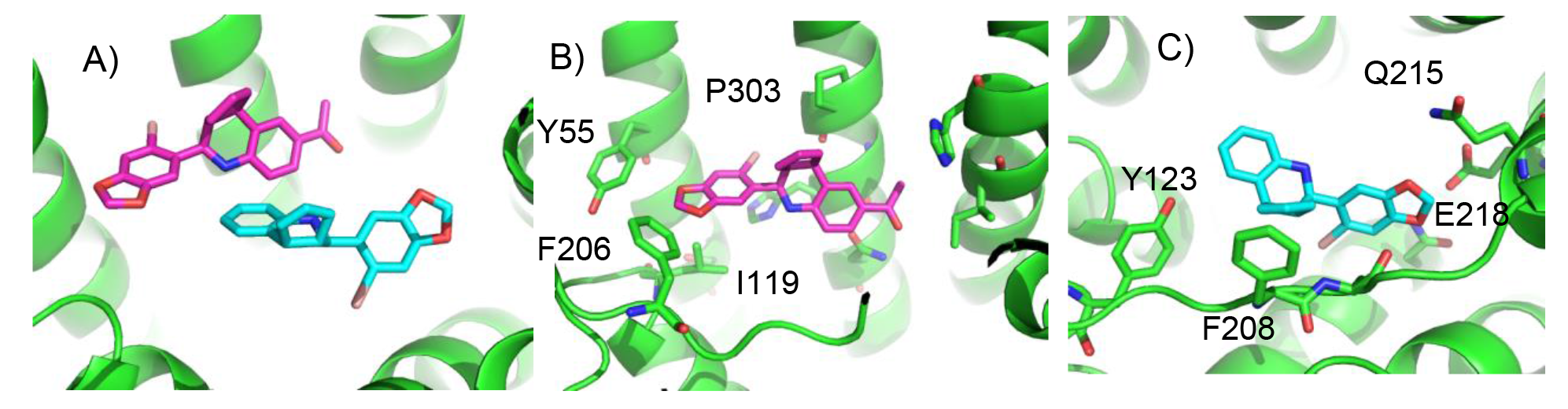
Predicted G1 and G15 binding sites in GPER. A) The highest scoring docking poses for G1 (maroon) and G15 (cyan). B) Receptor-ligand interactions for G1.C) Receptor-ligand interactions for G15.

ConDock predicted two equally high scoring, overlapping poses for tamoxifen, near E115, L119, Y123, L137, Q138, M141, Y142, Q215, E218, W272, E275, I279, P303, G306, H307, and N310 (Fig. 5). The conservation score of this orientation is 0.81. Hydrophobic interactions are observed between tamoxifen and non-polar residues L119, Y123, Y142, P303, and F314. Notably, the amine group of tamoxifen is neutralized by E218 and E275.

**Figure 5.**
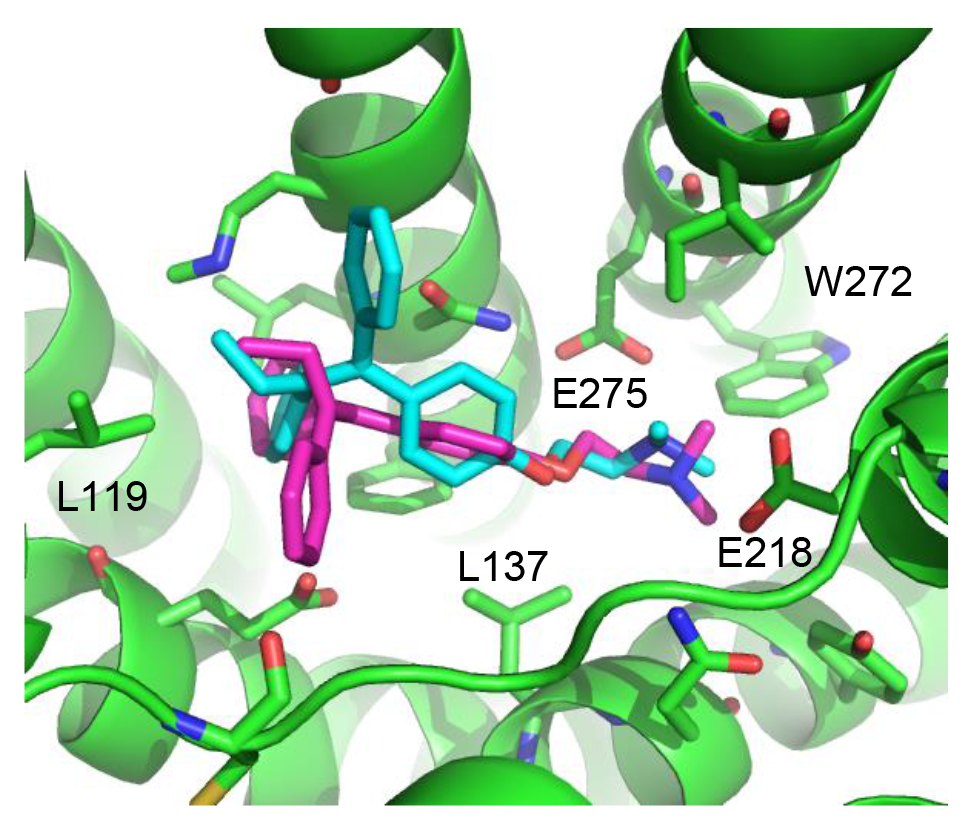
Predicted tamoxifen binding sites in GPER. A) The highest scoring docking poses for tamoxifen, pose 1 (maroon) and pose 2 (cyan).

In comparison with the ligand binding sites predicted by traditional methods based on surface geometry and conservation, the site predicted by ConDock is more detailed and of higher resolution due to the information from chemical interactions from ligand docking. Moreover, prediction methods based on surface geometry and conservation cannot differentiate between binding sites for different ligands. We compared the ligand binding site predicted by ConDock to those predicted by three other software packages representing different approaches: CASTp^46^, which analyzes surface geometry, SiteHound^47^, which maps surfaces with a chemical probe, and Concavity^33^, which analyzes surface geometry and conservation (Fig. 6). All three methods were able to identify a ligand binding site roughly matching that from ConDock. The pocket predicted by ConDock is deeper than the other pockets, which while intuitively attractive is not necessarily correct. SiteHound performed particularly poorly, with the top scoring site located on the GPER intracellular face. The site identified by SiteHound closest to the ConDock site was scored third and is a shallow binding pocket near H52-G58, E275-H282, and R299-H307 (Fig. 6C). In contrast, the Concavity site was smaller and more shallow than the ConDock site (Fig. 6D). Surprisingly, the site predicted by the simpler CASTp method best matched the ConDock site but is also smaller and more shallow (Fig. 6B). For proteins such as GPCRs with large, concave binding pockets, geometry based prediction methods such as Concavity and CASTp can easily identify the general location of the binding site. However, such methods may have more difficulty recovering the specific, ligand-specific binding site. It is also surprising that ConDock more closely matched the results of the geometry based methods given that ConDock does not take surface geometry into account. Without experimental data, it is not possible to conclude which of the predicted binding sites is correct at this time.

**Figure 6.**
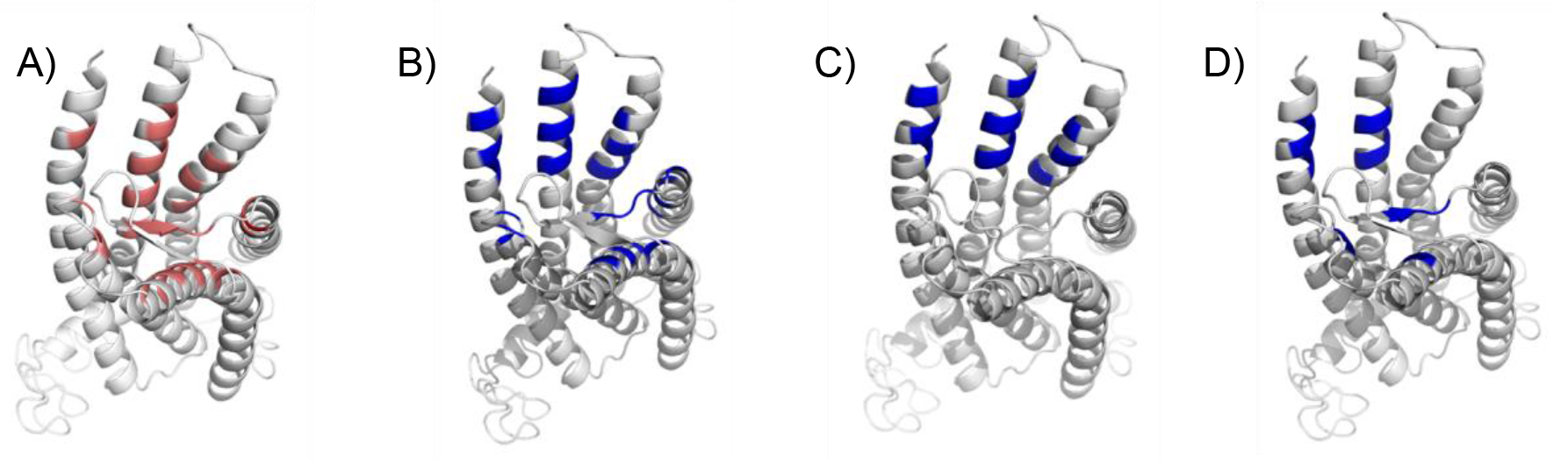
Predicted ligand binding sites by ConDock, CASTp, SiteHound, Concavity. Ligand binding sites are colored, predicted by A) ConDock, B) CASTp, C) SiteHound, D) Concavity.

## Discussion

The ConDock scoring method, incorporating information from both surface conservation and docking binding energy, was used to predict viable ligand binding sites for four different GPER ligands. In contrast to more typical geometry-based ligand binding site prediction methods, ConDock scoring takes advantage of chemistry-specific information about the ligand-receptor interface. The poor performance of SiteHound in predicting ligand binding sites on GPER suggests that a method based only on chemical interactions or docking is highly susceptible to error. Surface conservation data not only provides orthogonal knowledge but also dampens the influence from the shortcomings of current computational methods in homology modeling, docking, and predicting binding affinity. How best to mathematically combine these multiple data sources has been debated^29,33^, but we demonstrate here that a simple product function is effective. Although the ligands analyzed differ greatly in chemical structure, the ConDock scoring method predicted that all four bind to the same approximate region, deep in the extracellular cleft of the receptor. Undoubtedly, further refinement of a hybrid scoring function will lead to improved predictions.

Recent GPER modeling studies using molecular dynamics simulations and docking identified different potential binding sites for E2, G1, and G15 near F206 and F208; the interaction with this region was described as driven primarily by π-π stacking interactions^34,35^. Figure 7 compares the ConDock binding site against that predicted in the molecular dynamics simulation and docking study. The ConDock binding site is located deeper in the extracellular cleft; the other proposed site mostly involved surface exposed loops. Mendez-Luna *et al.* proposed that Q53, Q54, G58, C205, and H282 all interact with G1 and G15; however, none of these residues are conserved across the six species we analyzed. Experimental data is not currently available to support one model over the other.

**Figure 7.**
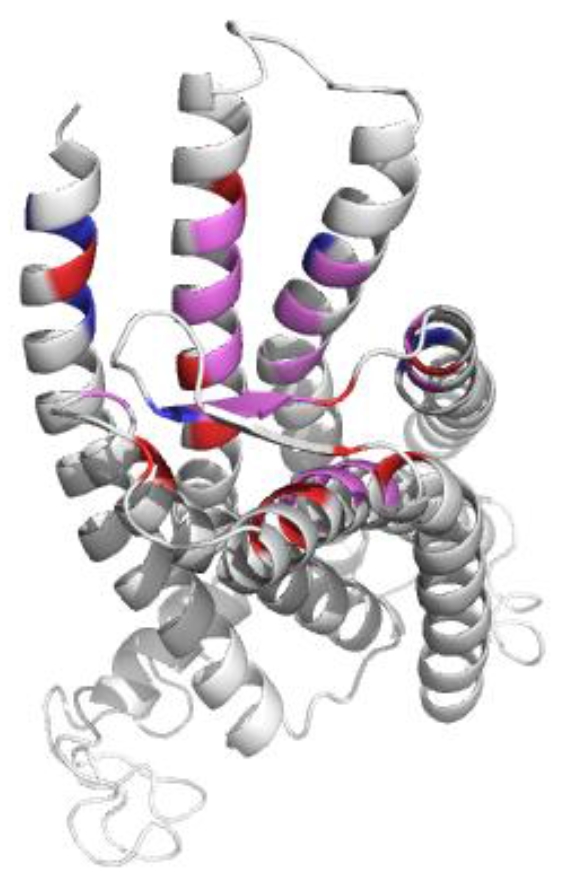
Comparison of proposed ligand binding sites. Comparison of the ConDock ligand binding site (red) with that proposed by Mendez-Luna *et al.* (blue). Residues found in both sites are colored violet.

In summary, the simple ConDock hybrid scoring model predicts physically plausible ligand binding sites by combining information from ligand docking and surface conservation. Using multiple orthogonal sources of information avoids errors introduced by modeling, especially in a case where a crystal structure of the receptor is unavailable. Using this hybrid method, we identified a site in the extracellular cleft of GPER that has the potential to bind four known GPER ligands. Further optimization of hybrid scoring functions should yield significantly improved predictions.

## Methods

### Protein conservation

GPER amino acid sequences were acquired from the UniProt protein sequence database^48^. Only Swiss-Prot^49^ curated sequences were compared using the Align function of UniProt. The amino acid sequences aligned were from human (*Homo sapiens*), rat (*Rattus norvegicus*), mouse (*Mus musculus*), Rhesus macaque (*Macaca mulatta*), zebrafish (*Danio rerio*), and Atlantic croaker (*Micropogonias undulatus*). The multiple sequence alignment file was submitted to ConSurf^43,44^. ConSurf assesses conservation using Bayesian reconstruction of a phylogenetic tree. Each sequence position is scored from 0-9, where 9 indicates that the amino acid was retained in all of the organisms (Fig. S4). Values from ConSurf were mapped onto the receptor surface with Chimera^50^.

### Homology modeling and docking

The crystal structure of GPER has not yet been determined. We created a model using GPCR I-TASSER (Iterative Threading Assembly Refinement), among the most accurate homology modeling softwares customized for GPCRs^36^. GPCR I-TASSER modeled the GPER structure using templates from the ten closest related GPCR crystal structures (PDB 4mbs, 2ks9, 1kpn, 1u19, 2ziy, 1kp1, 3odu, 4ea3, 4iaq, 2y00). The homology model was validated with ERRAT^37^. Coordinates for E2, G1, G15, and tamoxifen were downloaded from the ZINC ligand database^51^ and submitted to SwissDock^38^ for docking. SwissDock is a web interface to the EADock DSS^45^ engine, which performs blind, global (does not require targeting of a particular surface) docking using the physics based CHARMM22 force field^52^. The “FullFitness Score” calculated by SwissDock using clustering and the FACTS implicit solvent model^53^ was used as the “Energy Score” for our calculations.

### Combined analysis

SwissDock poses were manually screened for those binding sites located on or near the extracellular side of the protein. Ligand binding surfaces included residues within 3.5 Å from the docked ligand. The average conservation score of the amino acids that were highlighted served as the “Conservation Score” of that specific orientation (Fig. 8). The combined ConDock score is defined as the product of the Conservation and Energy Scores. As the Energy Score is a modified free energy function, a highly negative ConDock score is associated with a more probable ligand binding site. Binding sites predicted by ConDock results were compared with those predicted by CASTp^46^, SiteHound^47^, and Concavity^33^. For CASTp, SiteHound, and Concavity, ligand binding pockets were defined as residues within 4 Å of the selected probe/cluster.

**Figure 8.**
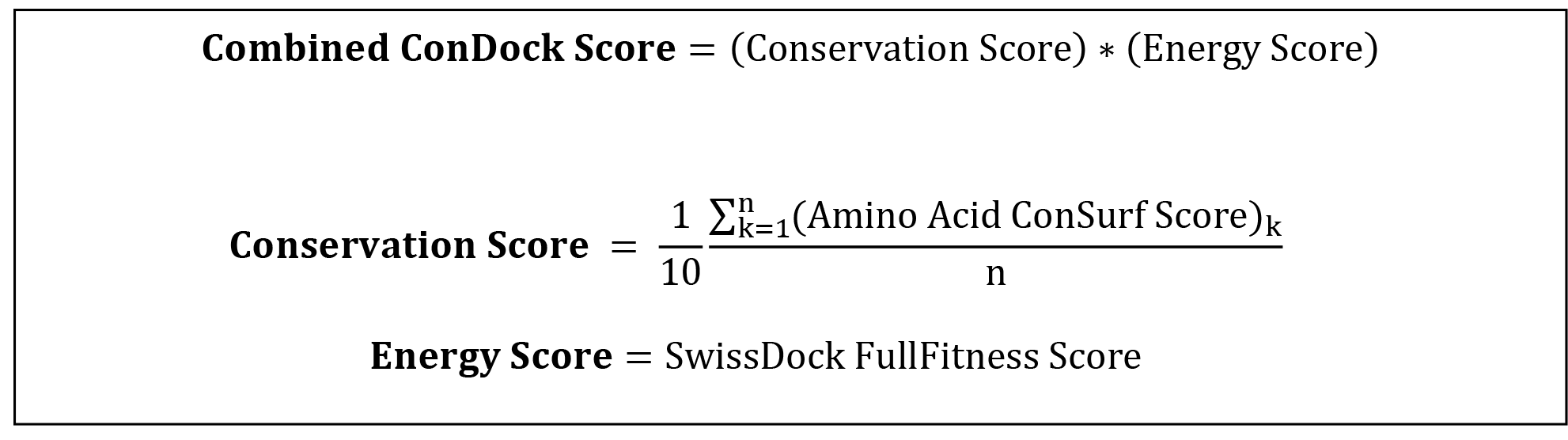
Calculation of combined ConDock scores for ligand binding sites. The Conservation Score is calculated over the n residues in a binding site, indexed by *k*.

## Acknowledgements

This work was funded by the Victoria S. and Bradley L. Geist Foundation (H.L.N.) CAREER Award 1350555 (H.L.N.), and the Undergraduate Research Opportunities Progr the University of Hawaii at Manoa (A.R.V., S.M.).

### Author contributions

A.R.V., S.M., and H.L.N. performed the analysis and calculations. A.R.V., S.M., and H. L.N. wrote the manuscript. H.L.N. supervised the project.

### Competing financial interests

The authors have no competing financial interests.

